# Standard melanoma-associated markers do not identify the MM127 metastatic melanoma cell line

**DOI:** 10.1101/026633

**Authors:** Parvathi Haridas, Jacqui A. McGovern, Ahishek S. Kashyap, D.L. Sean McElwain, Matthew Simpson

## Abstract

Reliable identification of different melanoma cell lines is important for many aspects of melanoma research. Common markers used to identify melanoma cell lines include: S100; HMB-45; and Melan-A. We explore the expression of these three markers in four different melanoma cell lines: WM35; WM793; SK-MEL-28; and MM127. The expression of these markers is examined at both the mRNA and protein level. Our results show that the metastatic cell line, MM127, cannot be detected using any of the commonly used melanoma-associated markers. This implies that it would be very difficult to identify this particular cell line in a heterogeneous sample, and as a result this cell line should be used with care.

## Introduction

Melanoma is an aggressive form of skin cancer that has the highest incidence rate in Australia^1^. Since many aspects of melanoma research rely on the use of various types of melanoma cell lines^2,3^, the reliable identification of different melanoma cell lines is very important.

A range of melanoma-associated markers are used to identify different types of melanoma cell lines^4^. The three most frequently used markers are: S100; HMB-45 and Melan-A^5, 6^. A common feature of many experimental investigations is that some melanoma cell lines are unable to be detected using certain markers^7^. To address this limitation, many studies use two different markers to ensure reliable identification^8^.

MM127 is a metastatic melanoma cell line of human origin, isolated in 1970^9^. Since then, MM127 melanoma cells have been used in many published investigations. For example, MM127 cells have been used in ultraviolet radiation studies^10^, gene-based studies^11^, drug-response studies^12^ and in other melanoma-associated research.

Some of our previous work involves investigating how the balance of the rate of cell migration and the rate of cell proliferation affects the collective spreading of a population of MM127 melanoma cells^13^. In this previous study we describe results from an *in vitro* monoculture circular barrier assay^14,15^, and we use a discrete random walk mathematical model to quantify the rates of cell migration and cell proliferation in the experiment. Because this previous study involves a monoculture assay with just one cell type present, we did not attempt to identify the MM127 cells using any melanoma-associated markers. One way to extend this previous work would be to perform more complicated co-culture experiments. Such an extension would require the identification of the MM127 melanoma cells amongst the total population of cells in the assay. To meet this aim we first need to establish whether we can reliably identify MM127 melanoma cells using standard melanoma-associated markers. The focus of the present study is to explore whether MM127 cells can be reliably identified using standard approaches.

This work is organised in the following way. In the Results section we describe the outcomes of three different experimental techniques for identifying different melanoma cell lines using S100, HMB-45, and Melan-A. These techniques include immunofluorescence, Western blotting, and quantitative reverse transcription-polymerase chain reaction (qRT-PCR) assays. These techniques are applied to four different melanoma cell lines, and the results for the metastatic melanoma cell line, MM127, are not as expected. We find that this cell line is not identifiable using any of the three markers.

## Results

### Short tandem repeat profiling

All melanoma cell lines: WM35; WM793; SK-MEL-28 and MM127, are validated using short tandem repeat (STR) profiling (Cell Bank, Australia. January 2015). The STR profiling results confirm that all these melanoma cell lines are identical to the reference samples held at Cell Bank (Supplementary information). Therefore, all melanoma cell lines in this study are identical to the reference samples held at Cell Bank.

The alleles obtained from STR profiling are analysed using the DMSZ database (http://www.dsmz.de/fp/cgi-bin/str.html) to give the closest match to each cell line we consider. The results from this analysis confirm that WM35, WM793 and SK-MEL-28 are as expected (Supplementary information). Conversely, the results for MM127 are not as expected since there is no match identified using the MM127 alleles [Fig. 1(a)]. This preliminary result suggests that further investigation is warranted.

**Figure 1:**
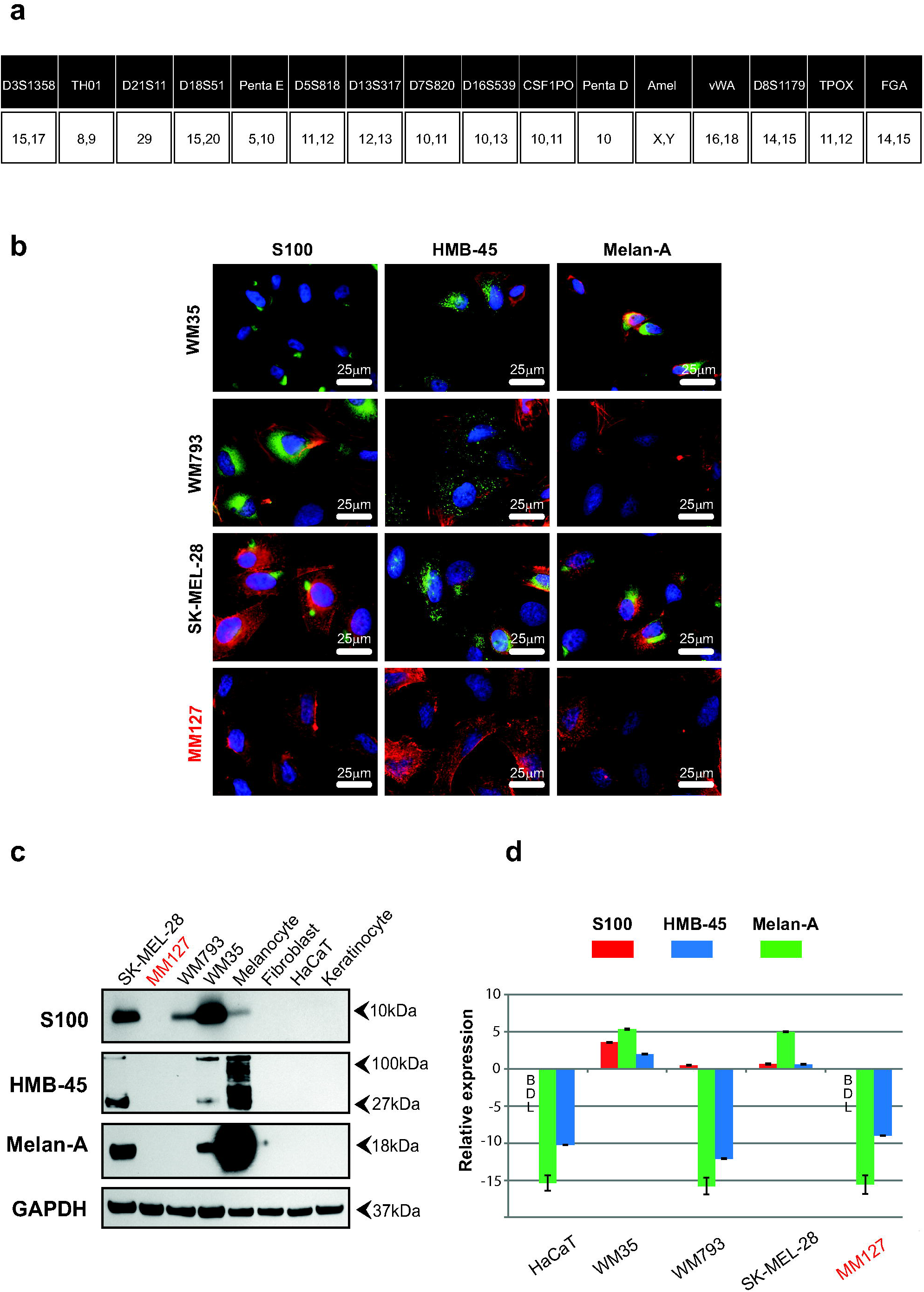
MM127 cell line does not express three standard melanoma-associated markers. (a) STR analysis of the MM127 cell line. Upper row shows the allele names. Lower row describes the location of the allele on the chromosome. (b) Immunofluorescence results for four melanoma cell lines (WM35, WM793, SK-MEL-28, MM127). Cells are fixed in 4% paraformaldehyde and stained for S100 (green), HMB-45 (green) and Melan-A (green). The nucleus (blue) and f-actin (red) are highlighted. Results for the negative control (HaCaT) are given in the supplementary material [Fig. S1]. (c) Melanoma (WM35, WM793, SK-MEL-28 and MM127) and negative controls (primary cells: keratinocytes, fibroblasts, and cell line: HaCaT) are analysed by Western blotting for S100, HMB-45 and Melan-A. GAPDH is used as loading control and detected at 37kDa. (d) qRT-PCR results show the difference between the expression of melanoma-associated genes for the negative control (HaCaT) and various cell lines. Values correspond to the mean ΔCt (n=3), where ΔCt = Ct (*RPL32*) – Ct (*target gene*). Error bars indicate the standard error (n=3). Results below the detectable limit are given as BDL.

### Immunofluorescence

Immunofluorescence is used on fixed cell preparations for S100, HMB-45 and Melan-A. The cell nuclei, f-actin and three standard melanoma-associated markers are highlighted [Fig. 1(b)]. S100 is localised to the nucleus and cytoplasm, and is observed in WM35, WM793 and SK-MEL-28 cells. HMB-45 is detected in the cytoplasm of WM35, WM793 and SK-MEL-28 cells [Fig. 1(b)]. Melan-A staining is present in both the WM35 and SK-MEL-28 cells, whereas it is absent from WM793 cells. Interestingly, MM127 is the only melanoma cell line that is negative for all three melanoma-associated markers in the immunofluorescence investigations. To confirm the immunofluorescence results, Western blotting analysis is performed.

### Western blotting

The expression of S100 (10kDa), HMB-45 (27 and 100kDa), and Melan-A (18kDa) proteins in the WM35, WM793, SK-MEL-28, MM127 and HaCaT cell lines are analysed using Western blots. The expression of these markers is also examined in primary cells, including keratinocytes and fibroblasts, showing that both keratinocytes and fibroblasts are negative for S100, HMB-45 and Melan-A [Fig. 1(c)]. The expression of these markers is also examined in primary melanocytes, showing that they are positive for S100, HMB-45 and Melan-A [Fig. 1(c)].

In the SK-MEL-28 and WM35 cell lines, HMB-45 is detected as two bands, which is consistent with previous results^16^. The cell lines WM793 and MM127 are negative for Melan-A and HMB-45 [Fig. 1(c)]. The absence of HMB-45 in WM793 cells in the Western blots does not coincide with the immunofluorescence analysis [Fig. 1(b)]. This is an interesting result that has not been reported previously. It is possible that HMB-45 could be detected by prolonging the exposure of the blot because HMB-45 is observed in the immunofluorescence results [Fig. 1(b)]. Discrepancies in protein expression among individual melanoma cells have been previously reported^17^, and this is consistent with the results presented here, since some individual WM793 cells are positive for HMB-45 [Fig. 1(b)] while other individual WM793 cells are negative for HMB-45. However, most importantly for our work, the expression of all three markers are absent from the MM127 cells. The Western blotting results for the MM127 cell line concur with the immunofluorescence results [Fig. 1(b)] and suggest that the MM127 cell line does not express the same antigens as the other melanoma cell lines investigated. To provide additional confirmation of the immunofluorescence and Western blot data, qRT-PCR assays are also performed.

### Quantitative reverse transcription-polymerase chain reaction (qRT-PCR)

The presence of genes that encode specific proteins associated with melanoma cell lines are quantified using qRT-PCR. The HaCaT cell line is used as a negative control because this cell line does not express any melanoma-associated markers as observed in immunofluorescence [Fig. S1], Western blotting [Fig. 1(c)] and qRT-PCR [Fig. S2] experiments. All qRT-PCR results are reported relative to the housekeeping gene (*RPL32*).

The qRT-PCR results indicate that the expression of S100 (*S100B*), Melan-A (*MLANA*) and HMB-45 (*PMEL*) in MM127 is very similar to the negative control [Fig. 1(d)]. The gene level for S100 in the HaCaT and MM127 cell lines is below the detectable limit (BDL). Overall, these qRT-PCR results are consistent with the Western blotting and immunofluorescence analysis, and confirm that these three standard melanoma-associated markers are not detected in the MM127 cell line.

To further verify the qRT-PCR results, the gene expression of two other commonly-used melanoma markers: microphthalmia-associated transcription factor (*MITF*); and tyrosinase (*TYR*)^18^ are also examined. Again, the expression of *MITF* and *TYR* in the MM127 cells are very similar to the negative control (Supplementary information).

## Discussion

Our group, and several other research groups, have previously worked with the metastatic melanoma cell line, MM127^13, 19-21^. Our previous work involves performing and analysing two-dimensional barrier assays with MM127 cells in monoculture^13,21^. Since these previous studies involve a monoculture experiment, there was no need to identify the cells within the experiment since all cells are known to be MM127 cells. If these monoculture experiments are to be extended to co-culture conditions, with multiple cell types present, we would be interested in identifying the MM127 cells within the total heterogeneous population. The results of the present study indicate that making this kind of distinction using standard melanoma-associated markers would be very difficult because MM127 cells cannot be identified using S100, Melan-A, HMB-45, MITF or tyrosinase.

Immunofluorescence on WM35, WM793 and SK-MEL-28 show positive expression of all the melanoma–associated markers considered, while the negative control, HaCaT cells, and the metastatic melanoma cell line MM127 did not express any of the markers we consider. All antibodies are used at an optimal dilution which we determined using a series of sensitivity assays specifically optimised for each marker prior to proceeding with the experiments. The Western blot analysis also shows the presence of all markers in WM35 and SK-MEL-28, and S100 in WM793. In contrast, all melanoma-associated markers considered are absent in the negative control and the MM127 cell line.

We find that the WM793 cell line does not express all the melanoma markers considered. One possible explanation for a variation between the immunofluorescence and Western blot analysis for WM793 cells is that a longer period of time might be needed to expose the blot. However, using a longer period of time might cause an over expression of proteins in the other lanes. We are not concerned by this variation since WM793 cells reliably express S100, and so can be identified using this marker. On the contrary, MM127 cells cannot be identified using any of the commonly used markers we consider.

Collectively, our findings indicate that the standard melanoma-associated markers we consider are not detected in the MM127 cell line. Although the MM127 cell line has been used in several previous studies^13,19-21^, we find that this cell line cannot be detected using standard melanoma-associated markers. Therefore, we suggest that other metastatic melanoma cell lines that express standard markers, such as the SK-MEL-28 cell line, ought to be used in preference to the MM127 cell line.

Since the MM127 cells we use in this study and the MM127 cells available from Cell Bank Australia are sourced from the same institution, it is not surprising that the cell validation results confirm that the cells we use are 100% identical to those available from Cell Bank (Supplementary information). However, since the MM127 cell line was first discussed in 1979^9^, it is possible that the MM127 cells currently available from Cell Bank are somehow different to the cells originally reported in 1979. Given that MM127 cells are primarily in use at one research institution, it is impossible to for us to repeat our investigations using samples of MM127 cells sourced from multiple institutions.

Certain recent investigations have shown that MM127 cells are positive for the *NRAS* mutation which is consistent with the idea that these cells are melanoma^22-23^. Regardless of their *NRAS* status, our observation that MM127 cells cannot be detected using standard melanoma-associated markers is an important finding. Although our study explores the presence of five different melanoma-associated markers in MM127 cells, it is always possible to extend our work by repeating the immunofluorescence, Western blotting and qRT-PCR experiments with additional markers. For example, recently it has been suggested that SOX10 is a sensitive marker for melanoma cells^24^, and so it would be interesting to repeat our work using SOX10. We have chosen not to use SOX10 in the present work because other authors suggest that S100 is a very sensitive marker for melanoma cells^5^.

## Methods

### Cell culture

Melanoma cell culture: Melanoma cell lines WM35 and WM793 are cultured in MCDB 153 medium (Sigma Aldrich, Australia) containing 20% Leibovitz L-15 medium (Life Technologies, Australia), 4% Foetal Calf Serum (FCS) (Hyclone, Australia), 7.5% w/v sodium bicarbonate (Life Technologies), 5μg/ml insulin (Sigma Aldrich), 1.68mM calcium chloride, 50U/ml of penicillin and 50μg/ml of streptomycin (Life Technologies). Melanoma cell lines MM127 and SK-MEL-28 are maintained in RPMI1640 medium (Life Technologies) supplemented with 10% FCS, 2mM L-glutamine (Life Technologies), 23mM HEPES (Life Technologies), 50U/ml of penicillin and 50μg/ml of streptomycin. The HaCaT cell line is cultured in Dulbecco’s Modified Eagle’s Medium (DMEM) (Life Technologies) with 10% FCS and 50U/ml of penicillin and 50μg/ml of streptomycin. All cells are routinely screened for *mycoplasma*.

### Primary Cell Culture

Keratinocyte culture: Human keratinocytes are isolated from skin discards collected after abdominoplasty and breast reduction surgery. All skin collections are obtained with informed patient consent and ethics approval (institutional and hospital: QUT 3865H 200 4/46). Keratinocytes are collected from the underside of the epidermis and papillary side of the dermis from surgical skin discards that are placed, overnight, in 0.25% trypsin (Life Technologies) diluted in a 1:1 ratio with phosphate buffered saline (PBS) (Life Technologies). Keratinocytes are grown in a 2:1 ratio on an irradiated feeder layer of 3T3 (i3T3) in full Green’s medium containing DMEM with Ham’s F12 (Life Technologies) in a 3:1 v/v ratio, 10% FCS, 2mM L-glutamine, 50U/ml of penicillin, 50μg/ml of streptomycin, 180mM adenine (Sigma Aldrich), 1μg/ml insulin, 0.1μg/ml cholera toxin (Sigma Aldrich), 0.01% non-essential amino acid solution (Life Technologies), 5μg/ml transferrin (Sigma Aldrich), 0.2μM triiodothyronine (Sigma Aldrich), 0.4μg/ml hydrocortisone (Sigma Aldrich) and 10ng/ml human recombinant EGF (Life Technologies). Cells are cultured at 37°C, in 5% CO_2_ and 95% air.

Melanocyte culture: Melanocytes are grown from an epidermal cell suspension (keratinocyte cell suspension). Isolated epidermal cell suspensions are seeded into T25cm^2^ tissue culture flasks (Nunc®, Australia) with 254 Medium (Life Technologies) together with Human Melanocyte Growth Supplement (Life Technologies). Epidermal cells, at a density of 8x10^4^cells/cm^2^, are seeded and incubated at 37°C in 5% CO_2_ and 95% air. If the melanocyte culture is contaminated with fibroblasts it is treated with 100μg/ml of Geneticin^®^ (Life Technologies) for 2-3 days.

Fibroblast culture: The dermis, obtained after keratinocyte isolation, is finely minced and placed in a 0.05% collagenase A type I (Life Technologies) solution prepared in DMEM at 37°C, in 5% CO_2_ and 95% air for 24 hours. The dermal cell solution is centrifuged at 212*g* for 10 minutes and the cells are seeded into T75cm^2^ flasks (Nunc®) in DMEM with 10% FCS, 2mM L-glutamine, 50U/ml of penicillin and 50μg/ml of streptomycin at 37° C in 5% CO_2_ and 95% air.

### Immunofluorescence

Cells are grown on glass coverslips and fixed with 4% paraformaldehyde (Electron Microscopy Sciences, Australia) for 30 minutes at room temperature. Cells are permeabilised with 0.1% Triton-X in PBS for 10 minutes, washed with 0.5% w/v bovine serum albumin (BSA) (Life Technologies) in PBS and the non-specific binding sites are blocked using 0.5% BSA for 30 minutes. This is followed by the addition of primary antibody (S100-1:2500, HMB-45-1:100 and Melan-A-1:200) (Dako, Australia) on cells for an hour, and the secondary antibody (Alexa Fluor® 480-1:400 and Alexa Fluor 555®-1:400) (Life Technologies) for an hour. Before proceeding with our experiments, we perform a series of experiments to determine the optimal dilution for each antibody. The optimal dilution was determined by starting with the manufacturer’s recommended dilution and then increasing the dilution to find an optimum result. Cells are washed three times with 0.5% BSA, the nucleus is stained with DAPI – 1:1000 (Sigma Aldrich) and f-actin is stained with Alexa Fluor® 488-1:200 (Life Technologies) for 10 minutes. Coverslips are mounted on glass slides using ProLong® Gold Antifade mountant (Life Technologies).

### Western blotting

Melanoma cell lines (WM35, WM793, MM127, SK-MEL-28) and non-melanoma cells (HaCaT, fibroblasts, melanocytes) are lysed by adding lysis buffer containing 80% Radio Immuno-Precipitation Assay (RIPA) buffer (Thermo Fisher Scientific, Australia), 10% protease inhibitor (Roche Diagnostics, Australia) and 10% phosphatase inhibitor (Thermo Fisher Scientific). Cells are collected in Eppendorf tubes, vortexed every 5 minutes for half an hour and passed through a 27.5 gauge needle three to four times. Cell lysates of keratinocytes are obtained following removal of the feeder layer. The cell pellet is washed twice with PBS, and the cells are lysed by adding the lysis buffer. Cell lysates are centrifuged at 18000*g* for 15 minutes at 4°C. The proteins are separated using sodium dodecyl sulphate polyacrylamide gel electrophoresis (SDS-PAGE) and transferred onto nitrocellulose membranes (BioTrace®NT, Pall Corporation, USA). Membranes are blocked in Tris-buffered saline containing 0.05% Tween-20 (TBST) and 5% skim milk (TBST/milk 5%). All additional immunostaining washes are performed using TBST at room temperature. The corresponding primary antibody is used to incubate membranes overnight at 4°C; (S100-1:2000 (Dako), HMB-45-1:100 (Thermo Fisher Scientific) and Melan-A-1:1000 (Dako)) in TBST containing 5% BSA (TBST/BSA 5%), and with a secondary antibody, horseradish peroxidase-conjugated anti mouse (1:5000) (R&D Systems, USA); anti-rabbit (1:5000) (R&D Systems, USA); in TBST containing 5% skim milk for 1 hour. We use GAPDH (1:100) (Cell Signalling, USA) as an internal loading control. These optimal dilutions for each antibody are determined with a series of dilution assays. Membranes are washed five times in TBST and developed with enhanced chemiluminescence, ECL solution (GE Healthcare Life Sciences, Life Technologies). Images are compiled using Adobe Illustrator® and the auto colour balance is adjusted in all figures.

### Quantitative reverse transcription-polymerase chain reaction (qRT-PCR)

Total cellular RNA is extracted from melanoma (WM35, WM793, MM127, SK-MEL-28) and non-melanoma cell lines (HaCaT) using Trizol Reagent (Life Technologies) following the manufacturer’s protocol. One µg of RNA is reverse transcribed and cDNA is synthesised using SuperScript® III First-Strand (Life Technologies). The qRT-PCR is performed using an ABI Prism 7500 Sequence Detection System (Applied Biosystems, USA) and SYBR Green (Life Technologies). PCR cycle parameters are the same for all primers: 40 cycles of denaturing at 94°C for 30 seconds, annealing at 60°C for 40 seconds and extension at 72°C for 50 seconds. To quantify the expression of mRNA in each cell line the cycle threshold (Ct) value of each cell line is subtracted from the corresponding Ct value for the internal control, *RPL32*. The main results are presented here [Fig. 1(d)] and additional results are given in the supplementary material document (Supplementary information). Each qRT-PCR assay is performed in triplicate. Data are reported as the sample mean obtained by averaging over the three identically-prepared replicate ± the sample standard error.

## Acknowledgements

This work is supported by the Australian Research Council (FT130100148, DP140100249). We thank Mitchell Stark for providing the MM127 cell line, and Nikolas Haass for providing all other melanoma cell lines used in this work and for providing comments on a draft version of this manuscript. We also thank Rachael Murray, Joan Roehl and Dominic Guanzon for technical assistance.

## Author Contributions

P.H., J.A.Mc., A.S.K., D.L.S.Mc., M.J.S. conceived the experiments. P.H. conducted the experiments. P.H., J.A.Mc., A.S.K., D.L.S.Mc., M.J.S. analysed the results. All authors reviewed the manuscript.

## Additional Information

Competing Financial Interests

The authors declare no competing financial interests.

